# A viral SAVED protein with ring nuclease activity degrades the CRISPR second messenger cA_4_

**DOI:** 10.1101/2025.07.01.662501

**Authors:** Marta Orzechowski, Ville Hoikkala, Haotian Chi, Stephen McMahon, Tracey Gloster, Malcolm F White

## Abstract

Type III CRISPR systems typically generate cyclic oligoadenylate (cOA) second messengers such as cyclic tetra-adenylate (cA_4_) on detection of foreign RNA, activating ancillary effector proteins which elicit a diverse range of immune responses. The CalpLTS system elicits a transcriptional response to infection when CalpL binds cA_4_ in its SAVED (SMODS associated and fused to various effectors domain) sensor domain, resulting in filament formation and activation of the Lon protease domain, which cleaves the anti-Sigma factor CalpT, releasing the CalpS Sigma factor for transcriptional remodelling. Here, we show that thermophilic viruses have appropriated the SAVED domain of CalpL as an anti-CRISPR, AcrIII-2, which they use to degrade cA_4_. AcrIII-2 dimers sandwich cA_4_, degrading it in a shared active site to short linear products, using a mechanism highly reminiscent of CalpL. This results in inhibition of a range of cA_4_ activated effectors *in vitro*. This is the first example of a virally-encoded SAVED domain with ring nuclease activity, highlighting the complex interplay between viruses and cellular defences.

## Introduction

Phages, viruses that infect bacteria, have coevolved with their hosts for billions of years, resulting in the development of a diverse range of prokaryotic defence mechanisms [1]. These defences range from simple mutations in cell surface receptors to the more complex adaptive immunity provided by CRISPR-Cas systems. In recent years, there has been a surge in the discovery of new defence systems, alongside the identification of novel mechanisms within previously known systems [2, 3].

Immune systems that function via generation of cyclic nucleotide second messengers, exemplified by the eukaryotic cGAS-cGAMP-STING system [4], are also commonly used in prokaryotic anti-viral defence. The bacterial ancestor of cGAS, named CBASS (Cyclic Oligonucleotide-Based Anti-Phage Signaling System) generates a wide range of cyclic nucleotide signalling molecules, activating a diverse range of downstream effectors [5-7]. Furthermore, type III CRISPR systems sense invading RNA and activate a specialised polymerase domain in the Cas10 subunit, generating a range of cyclic oligoadenylate (cOA) signalling molecules [8, 9]. These second messengers bind to a wide range of auxiliary effector proteins to provide an anti-viral response [10]. The most common sensor domains are the CRISPR-associated Rossmann Fold (CARF) and the distantly related SAVED domain [11]. Type III CRISPR systems can generate large quantities of cOA molecules, amplifying the primary signal of foreign RNA and ensuring full activation of the ancillary effectors [12] [13]. On the other hand, these systems generally encode a means to deactivate themselves using enzymes known as ring nucleases [14], which include a diverse range of stand-alone enzymes as well as the intrinsic ring nuclease activities of some effectors [15]. Virally encoded anti-defence proteins can intercept cyclic nucleotides to disable anti-viral defence systems. These include ring nucleases such as the anti-CRISPR AcrIII-1 [16] and cyclic nucleotide binding “sponges” such as the anti-CBASS proteins Acb1 and Acb2 [17, 18].

To expand the catalogue of viral-encoded type III anti-CRISPR proteins, we developed a simple bioinformatic search strategy, searching for short CARF or SAVED-containing proteins in phage genomes of the Millard database [19]. This search yielded a single candidate protein encoded by *Thermocrinus* Great Boiling Spring Virus (TGBSV) [20]. This 230 amino acid protein is closely related to the SAVED domain of the type III CRISPR protease CalpL. Two orthologues of CalpL have been studied previously, SsCalpL from *Sulfurihydrogenibium* spp [21, 22] and CcaCalpL from *Candidatus* Cloacimonas acidaminovorans [23]. Here, we demonstrate that the viral protein binds and degrades cA_4_, generating linear products, similar to the ring nuclease activity of the CalpL protein [22, 23]. The crystal structure reveals a dimeric organization consistent with “sandwiching” of cA_4_, which is cleaved using residues from both subunits. We tentatively name this protein AcrIII-2 (second anti-CRISPR of type III cA_4_-signalling defence systems) following the precedent set for AcrIII-1 [16].

## Results

### Identification of a candidate anti-CRISPR SAVED protein

In the past few years, six different families of stand-alone ring nucleases associated with type III CRISPR loci have been described [14-16, 24, 25]. In contrast, only one viral anti-CRISPR ring nuclease, AcrIII-1, has been identified [16]. To screen for new candidate viral ring nucleases, we developed a bioinformatic search strategy, searching for ORFs encoding CARF or SAVED domains in genomes of the Millard phage database [19]. This identified a promising candidate gene (NCBI accession: DAD54729) encoding a predicted viral SAVED protein (vSAVED) in the genome of the *Thermocrinus* Great Boiling Spring Virus (TGBSV). This 41 kb phage genome was assembled from metagenomic data and to date the virus has not been isolated or studied [20]. A spacer matching TGSBV was identified in the type III-A CRISPR locus of its likely host, *Thermocrinis jamiesonii*, a member of the phylum Aquificae with an optimal growth temperature around 80 °C [20]. A BLAST search against NCBI’s nonredundant protein database using vSAVED as query produced hits mostly against cellular CRISPR effector proteins with a C-terminal SAVED domain (Supplementary figure 1). The top hit, with 53% sequence identity, was to the type III-B CRISPR effector CalpL (Lon-SAVED) from *Thermocrinis jamiesonii*. CalpL uses its C-terminal SAVED domain to bind cA_4_, resulting in filamentation of CalpL and activation of the N-terminal Lon protease domain, which degrades an anti-Sigma factor to elicit a transcriptional response to phage infection [21, 23].

### vSAVED is a cA_4_-specific ring nuclease

To investigate the TGBSV vSAVED protein *in vitro*, we created a codon-optimised synthetic gene and cloned it into a plasmid for expression in *E. coli* with an N-terminal cleavable polyhistidine tag. The protein was expressed and purified to homogeneity (Supplementary figure 2). The ring nuclease activity of vSAVED was tested *in vitro* by incubating the protein with cA_4_ at a 1:10 ratio over 20 min at 60 °C and analysing reaction products by HPLC (Figure 1A). After 5 min, two smaller peaks flanking the cA_4_ peak appeared. These were identified via MS/MS as A_4_>P, linear tetra-adenylate with a cyclic phosphate at the 3’ terminus (longer retention time) and A_4_-P (shorter retention time). Within 10 min, the majority of cA_4_ had been converted to these cleavage products, with A_4_-P as the predominant species. At this time point minor levels of A_2_>P and A_2_-P additionally started appearing. After 20 min, A_2_-P predominated. This indicates that vSAVED degrades cA_4_ via a sequential cleavage pathway, generating the intermediates A_4_>P, A_4_-P, and A_2_>P, ultimately producing A_2_-P as the final degradation product (Figure 1b). The protein showed much weaker ring nuclease activity with cA_3_ and cA_6_ substrates (Supplementary Figure 3). Based on the ring nuclease activity of this viral protein, we propose the name AcrIII-2 (second anti-CRISPR of type III systems), based on the precedent of AcrIII-1 [16], as these enzymes have no plausible role other than inhibition of CRISPR defence and are unlikely to be specific for type III CRISPR subtypes.

**Figure 1.**
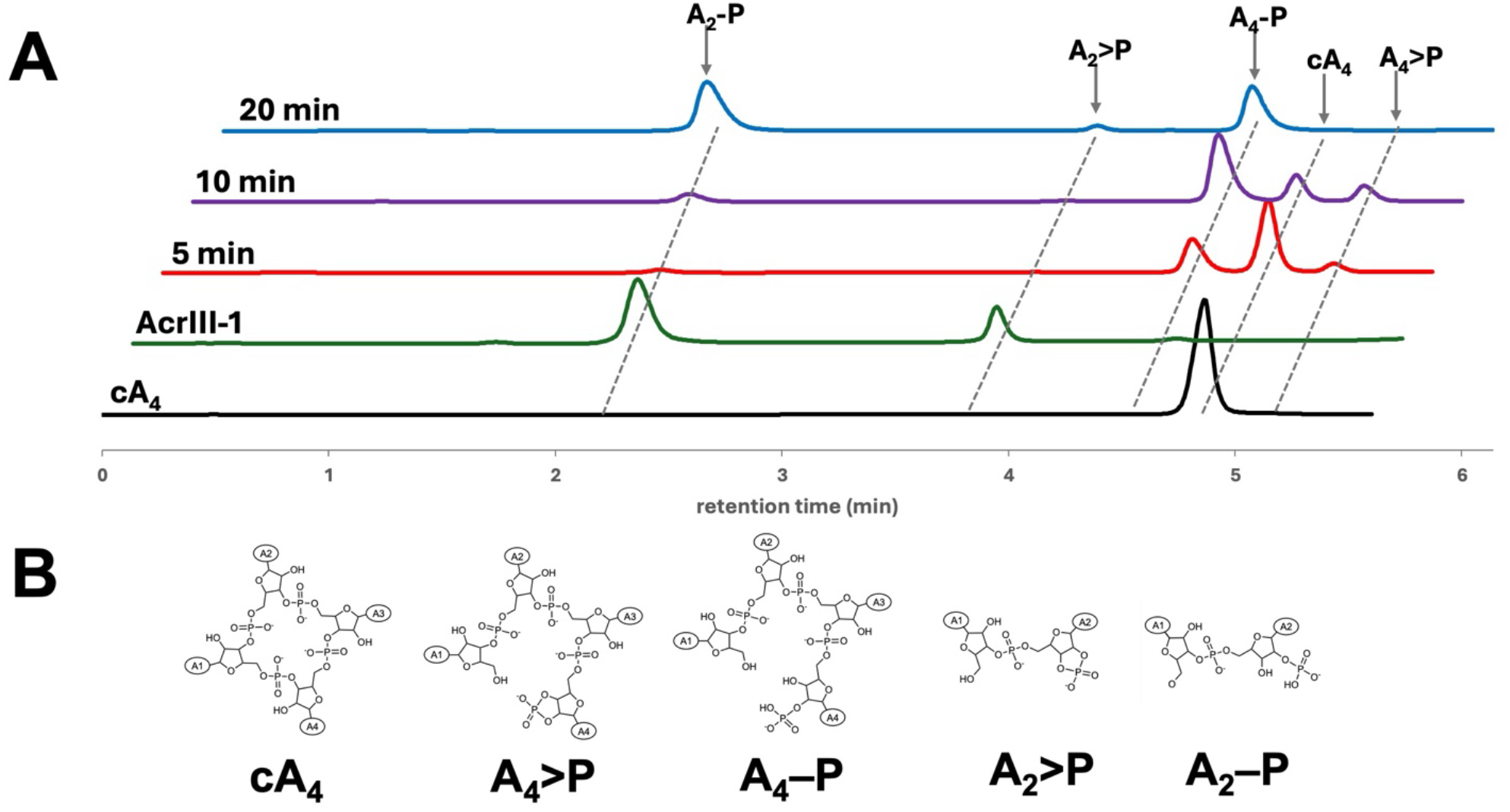
vSAVED (AcrIII-2) is a cA_4_-specific ring nuclease. **A**. HPLC traces of cA_4_ cleavage products when incubated with vSAVED for the indicated time periods. Reaction intermediates and products were identified using standards and/or mass spectrometry (Supplementary Figure 4). The AcrIII-1 ring nuclease, which generates A_2_>P and A_2_-P, was used as a control. **B**. Structural representation of cA_4_ and its cleavage products.

### AcrIII-2 inhibits cA_4_ activated effector proteins

We next tested whether AcrIII-2 could inhibit cA_4_-activated CRISPR effectors *in vitro*. We first examined TTHB144, a Csm6 family ribonuclease that is activated by cA_4_ [26]. In the presence of cA_4_, TTHB144 degrades RNA substrates, creating a measurable fluorescent signal in an RNase Alert assay [26]. In the assay, cA_4_ was used at a constant concentration (0.156 µM) that would allow for appropriate TTHB144 activation. AcrIII-2 levels were varied from 100 nM to 5µM during a 30 min preincubation step. At 250 nM AcrIII-2 (12X excess of cA_4_), enough of the signaling molecule remained in the assay to fully activate TTHB144 (Figure 2A). Once AcrIII-2 levels were above 500 nM (<= 6X excess of cA_4_), TTHB144 could no longer be activated, consistent with depletion of cA_4_ in the assay.

**Figure 2.**
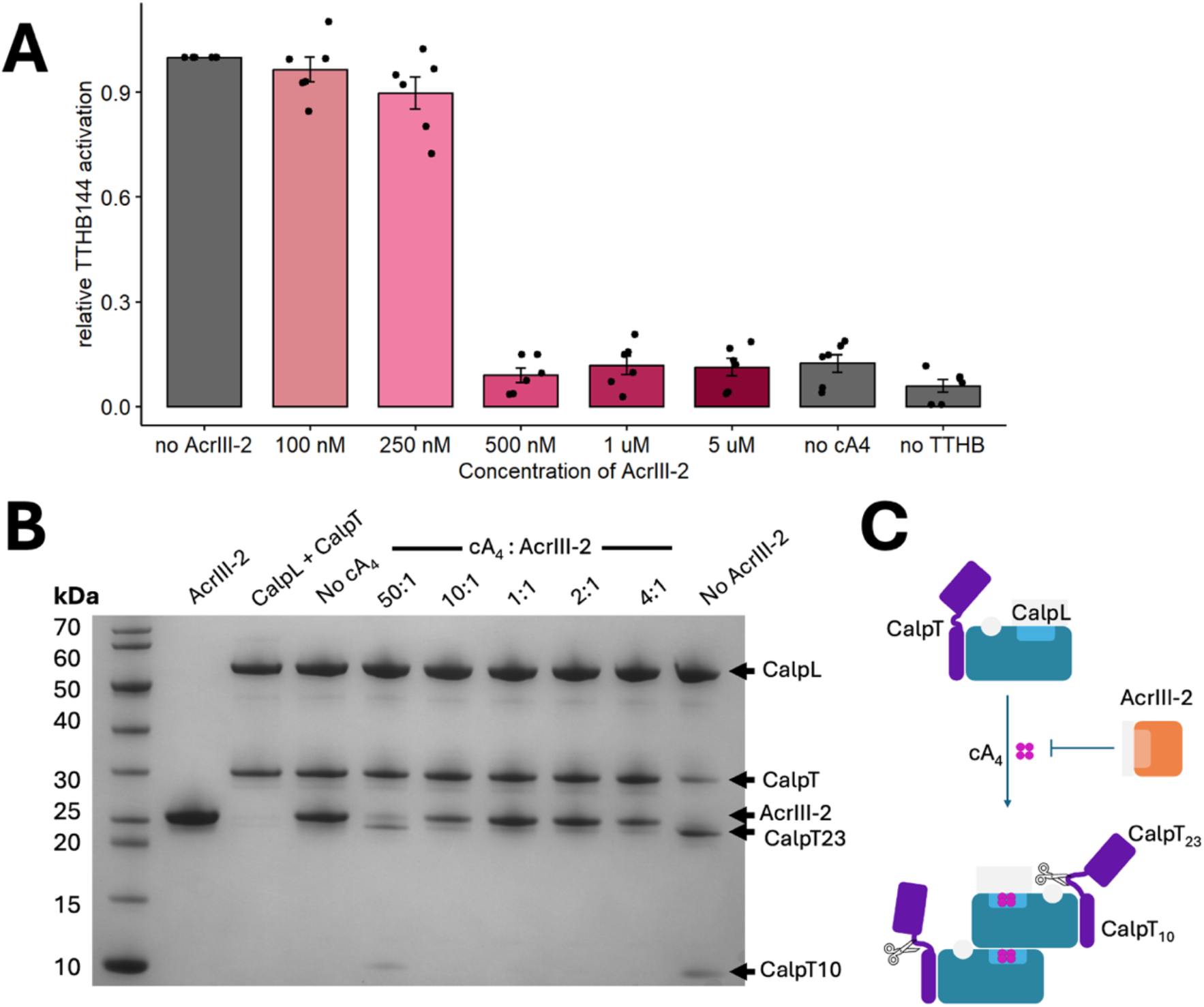
AcrIII-2 inhibits cA_4_specific effectors. **A**. AcrIII-2 limits activation of the cA_4_ activated ribonuclease TTHB144 at sub-stoichiometric ratios of AcrIII-2:cA_4_. Data are reported as mean ± SD of 6 technical replicates normalised to no AcrIII-2 control. **B**. AcrIII-2 can inhibit CalpT cleavage by SsCalpL. Cleavage products were visualised on an SDS-PAGE gel. When cA_4_ was preincubated with AcrIII-2, CalpL was only activated at a 50:1 ratio of cA_4_ to AcrIII-2, resulting in CalpT_23_ and CalpT_10_ bands appearing on the gel. This result is representative of an experiment done in triplicate. **C**. Illustration of the protease assay. Upon activation, Cas10 synthesises cA_4_ from ATP. The binding of cA_4_ by the preformed CalpL – CalpT complex results in oligomerisation and subsequent proteolytic cleavage of CalpT into CalpT_23_ and CalpT_10_. When cA_4_ is preincubated with AcrIII-2, this cleavage no longer occurs.

AcrIII-2 activity was further tested with the cA_4_ activated protease SsCalpL, which senses the signaling molecule through a SAVED domain [21]. Upon cA_4_ binding, SsCalpL triggers the cleavage of the associated protein CalpT into CalpT_23_ and CalpT_10_ (Figure 2B,C). To test the ability of AcrIII-2 to inhibit this cleavage mechanism, cA_4_ was preincubated with AcrIII-2 before being added to a mixture of CalpT and CalpL. The proteins and their cleavage products were then visualized on an SDS-PAGE gel (Figure 2B). The results show that in the presence of cA_4_, CalpT is cleaved into CalpT_23_ and CalpT_10_, seen by the disappearance of the CalpT band (around 30 kDa) concomitantly with the appearance of both a band at 23 kDa and 10 kDa. When no cA_4_ was added, this cleavage did not occur. The addition of just AcrIII-2, but no cA_4_, additionally did not affect the cleavage of CalpT. When AcrIII-2 was added to the assay at ratios of 4:1 to 1:10 with respect to cA_4_, no cleavage of CalpT could be observed. CalpL protease activity (appearance of the CalpT_23_ and CalpT_10_ bands) could be observed at a 1:50 ratio of AcrIII-2 to cA_4_, demonstrating that some uncleaved cA_4_ remained in the sample under these conditions.

Therefore, AcrIII-2 has the ability to inhibit cA_4_-activated CRISPR effectors at sub-stoichiometric concentrations with respect to cA_4_, consistent with the observed multiple turnover ring nuclease activity (Figure 1).

### AcrIII-2 multimerises in solution

To investigate the quaternary structure of AcrIII-2, we crosslinked the protein with bissulfosuccinimidyl suberate (BS3) in the presence or absence of cA_4_ and analysed crosslinked protein species by SDS-PAGE (Figure 3). The presence of dimeric crosslinked species was readily detected, along with small amounts of larger complexes. The extent of dimerisation was not significantly increased by the presence of cA_4_.

**Figure 3.**
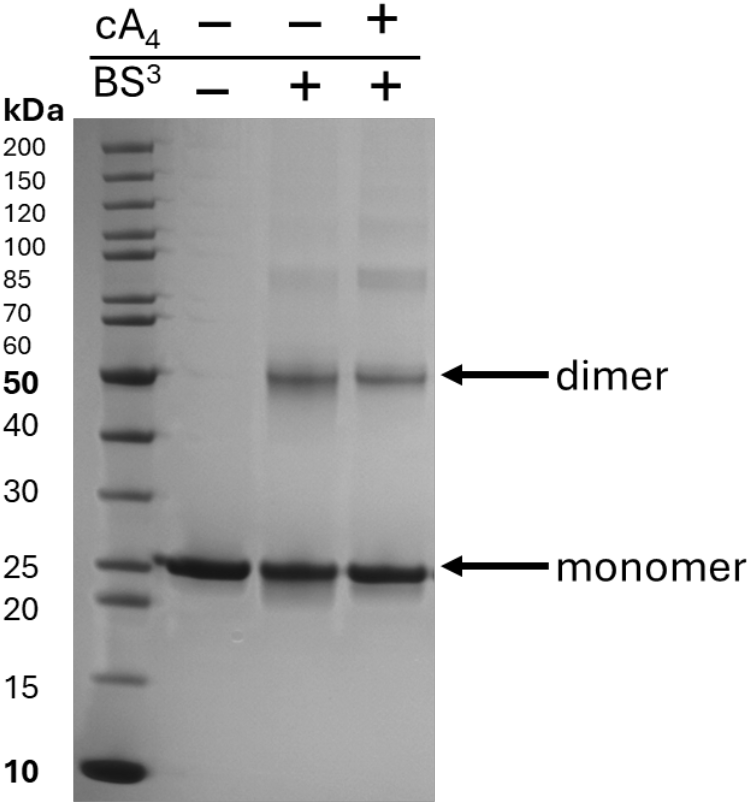
Crosslinking of AcrIII-2 with BS3. The addition of BS3 resulted in the formation of dimeric species and some larger oligomers of AcrIII-2. This was independent of the presence or absence of cA_4_. The gel is representative of experiments completed in triplicate.

### Structure of AcrIII-2

The structure of AcrIII-2, solved by molecular replacement with X-ray crystallography with data to 1.79 Å resolution, revealed two molecules in the asymmetric unit. Each molecule resembles a canonical SAVED subunit, which sit in a head-to-tail configuration (Figure 4A). The structure of an AcrIII-2 monomer comprises two CARF-like domains tethered by a 19-residue loop. Each CARF domain contains a five stranded β-sheet sandwiched between three α-helices; strands 1 to 4 of the β-sheet lie parallel, whilst the fifth strand is anti-parallel. The loop fusing the domains is followed by a 6-residue β-strand that lies anti-parallel to the first β-strand of the canonical C-terminal CARF domain, in essence making this a 6-stranded β-sheet. The AcrIII-2 SAVED domain monomer structure is a close match to CcaCalpL (PDB: 9EYJ) and SsCalpL (PDB: 8B0R) SAVED domains, with an RSMD of 2.2 Å (over 226 Cα atoms) and 2.5 Å (over 223 Cα atoms), respectively. for single subunits respectively). It proved difficult to directly superimpose the dimeric organisation of AcrIII-2 with the SAVED domains of the CcaCalpL bound to cA_4_ (PDB: 9EYJ), where cA_4_ is sandwiched between the two monomers. However, superimposition of the SAVED domains of CcaCalpL onto a AcrIII-2 monomer (Figure 4B) illustrated that while a monomer of each overlapped well, the other monomer was slightly offset. Whether this is caused by a subtle difference between the two proteins, or by binding of cA_4_, is unclear. It does, however, demonstrate that the global orientation of the two AcrIII-2 SAVED domains are not significantly different to the SAVED domains of CcaCalpL, meaning AcrIII-2 is highly likely to bind cA_4_ at the dimer interface and comparison to the CcaCalpL structure will help to define the likely AcrIII-2 cA_4_ binding site.

**Figure 4.**
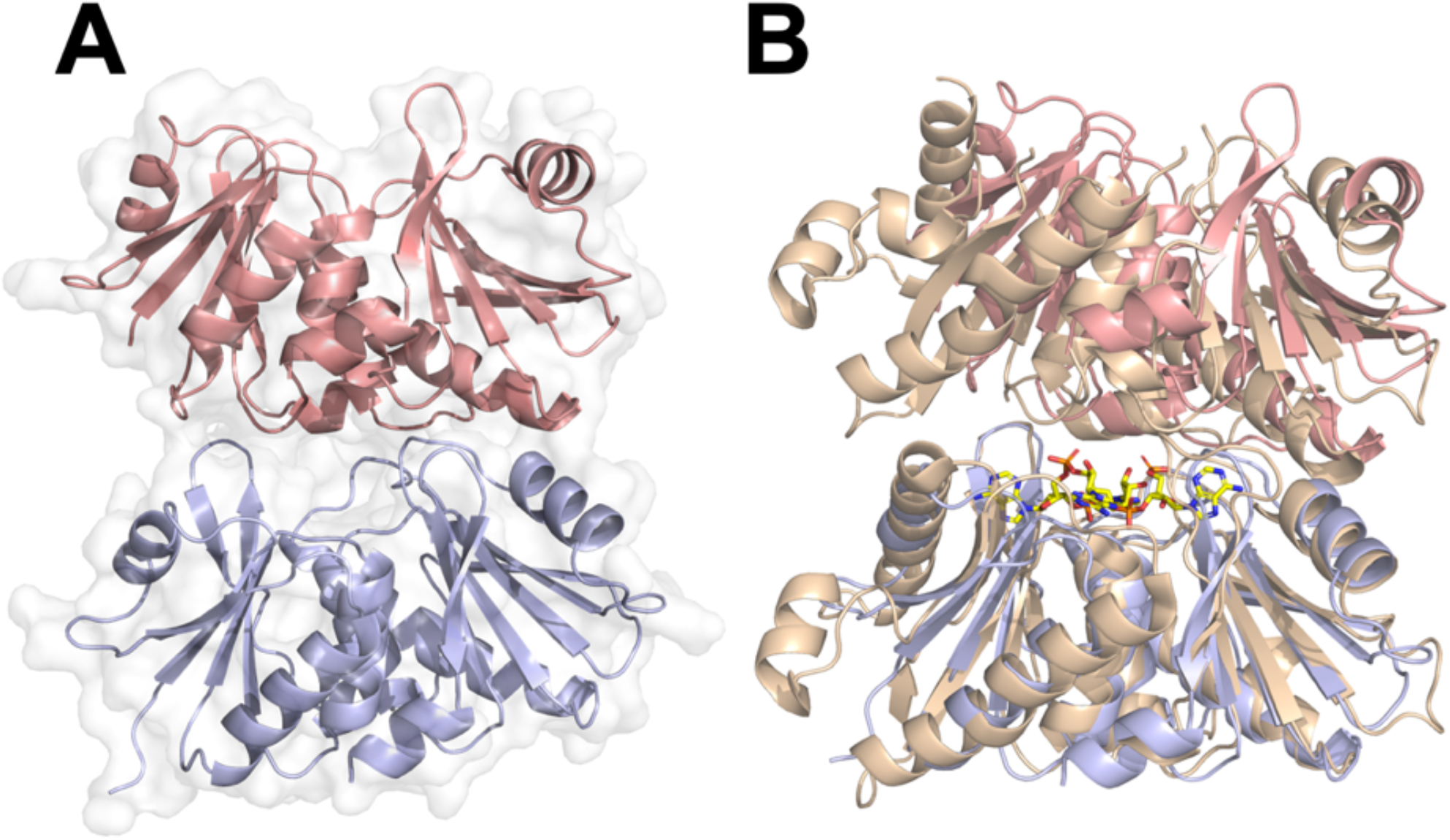
The structure of AcrIII-2 and comparison with CalpL. **A**. The structure of apo AcrIII-2. Each monomer of the dimer is shown in blue or salmon cartoon, with the grey outline showing the protein surface. **B**. Structural overlay of the AcrIII-2 dimer (blue and salmon cartoon) and the SAVED domains of CcaCalpL bound to cA_4_ (PDB: 9EYJ) (wheat) [23]. Note the alignment was based on a monomer (bottom one in the figure) of AcrIII-2 with the CcaCalpL dimer.

### AcrIII-2 dimers form a composite active site

Given the dimeric, head to tail structure of apoAcrIII-2 and the filament formation of CalpL when bound to cA_4_ [23], we hypothesized that AcrIII-2 dimers might use a composite active site for cA_4_ cleavage whereby residues from both monomers at the dimer interface contribute to cA_4_ binding and catalysis. To test this hypothesis, we identified potential residues from the AcrIII-2 structure that could fulfil important roles in cA_4_ cleavage, mutated these residues, and tested ring nuclease activity. In addition to H128 and S194 on the “lower” face of the binding site, we selected three conserved residues on the “upper” face (R96, K99, and Q178), generating the variants R96E, K99E, H128A, Q178A and S194A for analysis. All five variants were purified to homogeneity as for the wild-type protein (Supplementary Figure 2). Each variant was incubated with a 10-fold molar excess of cA_4_ for 30 min followed by separation of reaction products by HPLC (Figure 5).

**Figure 5.**
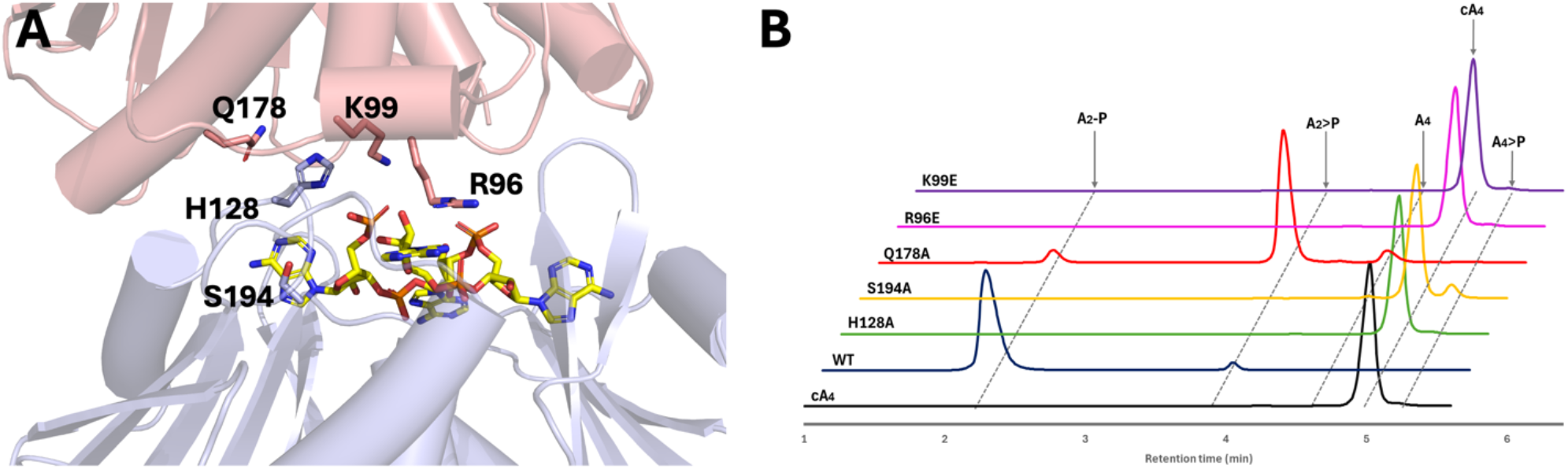
Mutational analysis of active site residues in the AcrIII-2 dimer. **A**. AcrIII-2 dimer (cartoon representation, with one monomer in salmon and the other in blue) with conserved residues selected for further analysis indicated. The cA_4_ molecule here (yellow sticks) is modelled using the structure of the SsCalpL monomer [21]. **B**. HPLC traces of cA_4_ cleavage products when incubated with AcrIII-2 variants at a 10:1 ratio of cA_4_ to AcrIII-2 for 30 min, with a final cA_4_ concentration of 50 µM.

The results revealed the complete abolition of ring nuclease activity for the R96E, K99E and H128A variants. The S194A variant showed only very minimal conversion of cA_4_ to A_4_>P. The Q178A variant degraded cA_4_ to A_2_>P, with only minor levels of A_2_-P produced, suggesting a role for this residue in the hydrolysis of the cyclic phosphate moiety in the linear dinucleotide products following cleavage of cA_4_. When the concentration of AcrIII-2 was increased ten-fold to reach a 1:1 ratio of cA_4_ to AcrIII-2, the reaction products for the R96E, H128A and Q178A variants were unchanged, whilst K99E generated minor levels of A_4_>P. The S194A variant could degrade all the cA_4_ to linear tetra-adenylates under these conditions (Supplementary figure 5).

These observations indicate the importance of residues on both faces of the dimeric interface for cA_4_ degradation, confirming the existence of a composite active site for AcrIII-2. This corresponds to the situation for CcaCalpL [16], where H396 and R358, equivalent to H128 and R96 in AcrIII-2, are essential for ring nuclease activity, while K361 (equating to K99) plays a subsidiary role [23]. Similar results were obtained for SsCalpL [22]. Q178 in AcrIII-2 corresponds to N455 in CcaCalpL, which has not been studied [23].

### Inactive AcrIII-2 variants can be combined to rescue ring nuclease activity

A diagnostic test for proteins that share an active site across a subunit interface is to combine two inactive variants and test for the recovery of enzyme activity [24, 27]. We first tested whether AcrIII-2 and its active site variants multimerized upon cA_4_ binding using dynamic light scattering (DLS). Addition of cA_4_ to wild-type AcrIII-2 resulted in a significant increase in the average particle size (Figure 6A). This was also observed for the H128A and S194A variants, suggesting that these variants retain the ability to bind cA_4_ and multimerize. Although the average particle size was notably smaller, the R96E and K99E variants also displayed multimerization upon addition of cA_4_ (Figure 6A), suggesting that these charge reversal mutations may have weakened, but not abolished, the inter-subunit interface. Indeed, a double mutant with both R96E and K99E mutations also retained the ability to multimerize on cA_4_ binding. In contrast, the equivalent double mutation in CcaCalpL abolished filament formation [23]. Finally, mixing of equal amounts of the H128A and R96E variants yielded an increase in average particle size upon addition of cA_4_, broadly equivalent to that of R96E alone.

**Figure 6.**
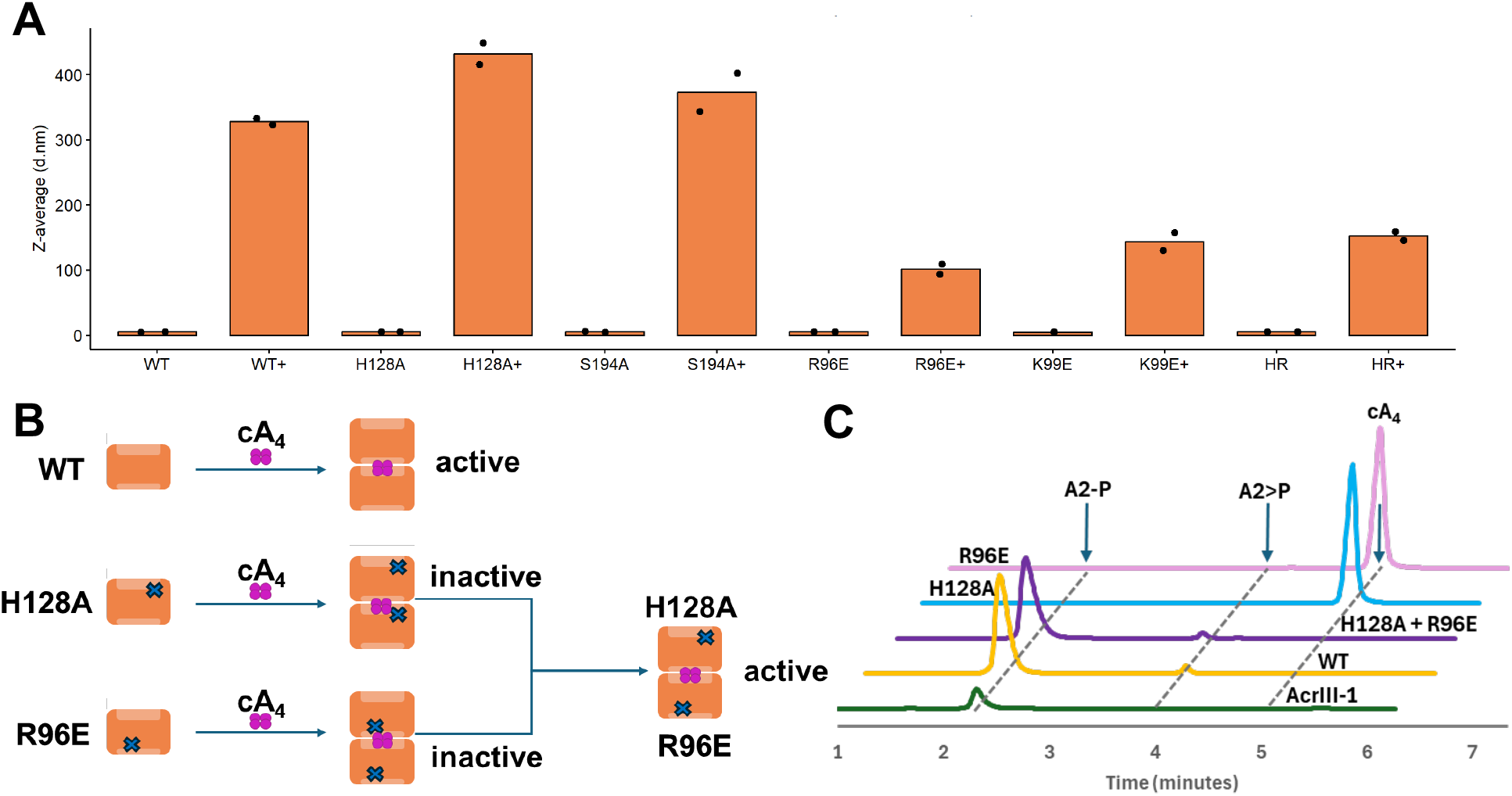
AcrIII-2 dimerises to degrade cA_4_. **a**. Z-average of AcrIII-2 WT and variants at a sample concentration of 1 mg/mL (37 μM) in the presence and absence of 10X cA_4_ determined using DLS. Samples were measured in duplicates and individual data points are shown. **B**. Schematic representation of subunit mixing assay. **C**. HPLC analysis of cA_4_ cleavage products after 30 min for individual and mixed variants. The cleavage products of ring nuclease AcrIII-1 are shown as a standard for A_2_-P.

Given the ability of AcrIII-2 variants to multimerize upon addition of cA_4_, we proceeded to investigate ring nuclease activity when combining the two inactive variants H128A and R96E, which have the potential to form a “wild-type” active site upon dimerization (Figure 6B). Although inactive in isolation, the combination of these two variants allowed recovery of ring nuclease activity for AcrIII-2 (Figure 6C). These data provide unequivocal confirmation that AcrIII-2 functions via a composite active site that forms between subunits, sandwiching the cA_4_ substrate. Similar results were recently obtained for the CalpL proteins [22, 23].

## Discussion

The virally-encoded SAVED protein studied here has all of the properties expected of an anti-CRISPR protein, and is thus named AcrIII-2 following the precedent set by AcrIII-1 [16]. As the first example of an Acr based on a SAVED domain, AcrIII-2 represents a clear case of “genetic scavenging” of a bacterial defence protein for use in an anti-defence role. The crystal structure of AcrIII-2 revealed a dimeric composition in the absence of cA_4_, with SAVED subunits orientated similarly to that observed in the cryoEM structure of the CalpL filament bound to cA_4_ [23]. Coupled with the observation of a dimeric species formed by cross-linking in the absence of cA_4_, these data suggest that apo AcrIII-2 dimerises, and it is likely the dimers and longer filaments are stabilized upon cA_4_ binding.

AcrIII-2 is clearly related to the SAVED domain of the CalpL (Lon-SAVED) protein, which is itself a ring nuclease [22, 23]. Key catalytic residues are conserved, and the observation of step-wise conversion from cA_4_ to linear A_4_ and hence A_2_ products is a hallmark of both proteins. This proposed mechanism requires release of A_4_ and rebinding in a suitable position for a second round of cleavage, as posited for CalpL [23]. Despite the similarities of AcrIII-2 with the SAVED domain of CalpL, its primary role may not be to prevent activation of CalpL. In fact, CalpL is very rarely found as a sole effector in type III CRISPR systems and is commonly present in type III CRISPR loci in genomes of thermophiles along with a second cA_4_-activated effector such as Csx1, Can2 or Cam1 [10]. Effector combinations are observed frequently in type III CRISPR loci, and given the role of the CalpL system in transcriptional reprogramming it is perhaps not surprising that it is often coupled with a more canonical effector that degrades nucleic acids [28-30] or collapses the membrane gradient [31]. Notably, Csx1, Can2 and Cam1 often do not possess intrinsic ring nuclease activity [15], so a virally-encoded ring nuclease like AcrIII-2 could be effective in the neutralization of type III CRISPR defence, even when the auto-deactivating CalpL system is present.

One important limitation of our study is that Acr activity has not been demonstrated *in vivo*. This is due to a technical hurdle, as AcrIII-2 is functional in very high temperature environments, making it more challenging to reconstitute in an *in vivo* experiment. Furthermore, the viruses and host strains necessary for these experiments have not yet been developed for lab studies. As such, the designation of the viral SAVED protein as an Acr, while highly plausible, is unproven.

Genes encoding SAVED domains that are not part of a larger effector are rare in bacterial genomes, but we identified two instances (WP_069292934 and WP_041081292) where a SAVED protein with conserved residues implicated in ring nuclease activity is observed as part of a type III CRISPR locus. In this context they presumably function as cellular ring nucleases, and we propose the name Crn5 (CRISPR ring nuclease 5) for this family, following the precedent whereby AcrIII-1 orthologues present in CRISPR loci are denoted “Crn2” [16].

## Methods

### Cloning

Cloning of the *acrIII-2* gene (Thermocrinis Great Boiling Spring virus; DAD54729.1) was completed according to standard protocols. Briefly, the synthetic gene (g-block) for AcrIII-2 was purchased from Integrated DNA Technologies (IDT) and ligated into the pEhisV5TEV vector between *Nco*I and *Bam*HI restriction sites [32]. The construct was transformed into competent DH5α *Escherichia coli* cells. The inclusion of the correct sequence was confirmed by sequencing (Eurofins Genomics). For protein expression, the plasmid was transformed into competent C43 (DE3) *E. coli* cells. Mutants were created by introducing point mutations using primer-directed mutagenesis into the plasmid.

### Protein expression

C43 (DE3) *E. coli* cells (Sigma Aldrich), transformed with the plasmid expressing the *acrIII-2* gene, were used for protein expression. A culture was grown in 2 L lysogeny broth (LB) at 37 °C with shaking at 180 RPM until an OD_600_ of 0.8 was reached. To induce protein expression, 0.4 mM isopropyl-β-D-1-thiogalactoside (IPTG) was added to the culture. Cells were then grown at 16 °C with shaking at 180 RPM overnight. The cells were harvested by centrifuging at 5000 RPM at 4 °C for 10 min (Beckman Coulter Avanti JXN-26, JLA-8.1000 rotor).

### Protein purification

For protein purification, the cell pellet was resuspended in 5X lysis buffer (50 mM Tris, 0.5 M NaCl, 10 mM imidazole, 10% glycerol, pH 7.5), with 1mg/mL lysozyme and 1 protease inhibitor tablet (Roche). The resuspended pellet was lysed by sonication on ice for six times 1 min with 1 min rest intervals. Cell debris was removed by ultracentrifugation of the lysate at 40,000 RPM for 30 min at 4 °C (Beckman Coulter Optima L-90K, 70Ti rotor).

AcrIII-2 was purified using a 5 mL HisTrap FF column (GE Healthcare), which was equilibrated using wash buffer (50 mM Tris-HCl, 0.5 M NaCl, 30 mM imidazole, 10% glycerol, pH 7.5). The supernatant collected after ultracentrifugation was loaded onto the column using an NGC chromatography system (Biorad) and unbound protein was eluted) using 20 column volumes (CV) of wash buffer, followed by a stringent wash using 10% elution buffer (50 mM Tris-HCl, 0.5 M NaCl, 0.5 M imidazole, 10% glycerol, pH7.5) for 3 CV. The His-tagged protein was eluted using a linear gradient of elution buffer across 15 CV and fractions analysed by SDS-PAGE.

Protein-containing fractions were concentrated and the 8-His affinity tag was removed by incubating the protein with Tobacco Etch Virus (TEV) protease (1 mg per 10 mg protein) overnight at room temperature while dialysing into wash buffer. Cleaved AcrIII-2 was further purified by repeating the immobilised metal affinity chromatography step and collecting the unbound fraction. Size exclusion chromatography (S200 16/60 column; GE Healthcare) was performed, with protein eluted using gel filtration buffer (20 mM Tris, 250 mM NaCl, 10% glycerol, pH 7.5). After SDS-PAGE to judge purity, relevant fractions were pooled, concentrated using a 10 kDa molecular weight cut-off centrifugal concentrator (Amicon Ultra-15), aliquoted, and frozen at -70 °C.

### Crosslinking

Purified protein was dialysed into X-link buffer containing 100 mM Na_2_PO_4_, 250 mM NaCl, and 10% glycerol, at pH 7.4 overnight. cA_4_ was used at a concentration 10-fold higher than the protein. Bis(sulfosuccinimidyl)suberate (BS3, Thermo Fisher) was used at a 1:5 molar ratio of protein:BS3. X-link buffer was used to adjust each reaction to 10 µL volume, and were incubated for 30 min at 20 °C. Reactions were subsequently quenched by addition of 2 µl 100 mM Tris pH 7.5 and analysed by SDS-PAGE.

#### Crystallisation

AcrIII-2 was concentrated to 35 mg mL^−1^ and centrifuged at 20000 x *g* prior to use. Sitting drop vapour diffusion experiments were set up at the nanoliter scale using commercial crystallisation screens and incubated at 293 K. The optimum crystals, as assessed by data quality, were obtained from 1.2 M sodium dihydrogen phosphate, 0.8 M dipotassium hydrogen phosphate, 0.2 M lithium sulphate, and 0.1 M CAPS pH 10. Crystals were cryoprotected with 25% glycerol prior to harvesting and cryo-cooled in liquid nitrogen prior to data collection.

### X-ray data collection and processing

Two X-ray data sets from the same AcrIII-2 crystal were collected at a wavelength of 0.9537 Å, 100 K, on beamline I04 at the Diamond Light Source. The data were processed and scaled together using xia2 multiplex [33]. The structure was solved using molecular replacement by PhaserMR [34] in the CCP4 suite [35] using a model generated by Alphafold2 [36] implemented in ColabFold [37], with initial B-factors modelled in Phenix [38]. Model refinement was achieved by iterative cycles of REFMAC5 [39] with manual model manipulation in COOT [40]. The quality of the structure was monitored throughout using Molprobity [41]. Data and refinement statistics are shown in Supplementary Table 1. The coordinates and data have been validated and deposited in the Protein Data Bank with deposition code 9R8S.

### TTHB144 ribonuclease assay

Assays were carried out in a buffer containing 20 mM Tris-HCl, 100 mM NaCl, pH 7.5. AcrIII-2 was incubated with cA_4_ at various ratios for 30 min at 60 °C. These reactants were then added to RNAse-Alert (IDT) substrate (final concentration 100 nM) and the *Thermus thermophilus* HB8 cA_4_ activated ribonuclease TTHB144 [26] (final concentration 125 nM), in a Greiner 96 half area plate. These reactions were incubated in a fluorescence plate reader (FLUOstar Omega) for up to 75 min at 45 °C with fluorescent measurements taken every 30 s (λex: 485 nm; λem: 520 nm).

### CalpL Protease assay

A buffer of 20 mM Tris, 50 mM NaCl, pH 8.0 was used to prepare all protein solutions. CalpL, a cA_4_-activated protease [21], was used at a final concentration of 1.73 μM. CalpT, a CalpL-associated protein, was used at a final concentration of 1.64 μM. Initially, cA_4_ was incubated with various concentrations of AcrIII-2 for 30 min at 60 °C. cA_4_ was used at a final concentration of 5 μM or 12.5 μM (for the 10:1 and 50:1 ratios respectively). Subsequently, CalpT and CalpL were added to the AcrIII-2 and cA_4_ reaction and incubated for another 30 min at 60 °C. The reaction was quenched by addition of SDS loading buffer and analysed by SDS-PAGE.

### HPLC

All proteins were prepared using a buffer of 20 mM Tris, 50 nM NaCl, and pH 8. cA_4_, cA_3_, cA_6_, and AcrIII-1 were used at a final concentration of 50 μM. AcrIII-2 was used at a final concentration of 5 μM. cA_4_ was incubated with AcrIII-2 for various time points at 60 °C. The reaction was stopped by the addition of 2.5X (v/v) methanol. cA_4_ and its cleavage products were extracted using methanol precipitation. Subsequent separation was performed using a Thermo Scientific Velos Pro Spectrometer equipped with a Dionex UltiMate 3000 chromatography system and a Kintex 2.6 μM EVO C18 column (Phenomenex, 2.1 ×100 mm). After injection of 5 μL sample, the components were eluted with solvent A (20 mM Ammonium acetate, pH 8.5) and solvent B (methanol) at a flow rate of 0.3 mL min^-1^ using a gradient of 0-3.5 min, 0% B; 3.5-5 min, 20% B; 5-7 min, 50% B; 7-10 min, 100% B. The column temperature was set to 40 °C and UV data were recorded at 260 nm.

### Mass spectrometry

Liquid chromatography tandem mass spectrometry (LC-MS/MS) was performed on a Eksigent 400 LC system coupled to a Sciex 6600 QTOF mass spectrometer, operated in trap-elute mode at microflow rates. Samples were injected into a YMC Triart C18 trap cartridge (0.5 × 5.0 mm) using 99.95% H_2_O with 0.05% TFA at a flow rate of 10 μl/min for 3 min to remove salts (diverted to waste). The trap was subsequently brought in-line with the analytical column (YMC Triart, 150 × 0.075 mm). Chromatographic separation was achieved using a gradient elution with solvent A (99.9% H_2_O with 0.1% formic acid) and solvent B (80% acetonitrile, 20% H_2_O and 0.1% formic acid) at a flow rate of 5 μl/min, following 0 min, 3% B; 0–6 min, 3-95% B; 6–8 min, hold at 95% B; 8–9 min, return to 3% B; 9–13 min, re-equilibrate at 3% B. The elute was directly introduced into the ESI turbospray of the mass spectrometer. Data were acquired in positive ion mode over amass range of m/z 120 – 1,000. Selected precursor ions were subjected to CID fragmentation using collision energies between 25-45 V and product ion spectra were collected.

### Dynamic Light Scattering (DLS)

The hydrodynamic radii of AcrIII-2 in the presence and absence of cA_4_ were measured by dynamic light scattering using a Zetasizer Nano S90 (Malvern) instrument. Samples with a final concentration of 37 μM (1mg/mL) AcrIII-2 were prepared in 20 mM Tris, 50 mM NaCl, pH 8 and both protein and cA_4_ stocks were centrifuged at 14,000 RPM for 10 min prior to use. A 10X concentration of cA_4_ was added to AcrIII-2 immediately prior to measurement. The experiment was carried out at 4 °C with 3 measurements of 13 runs. All experiments were carried out in at least duplicate.

## Data Availability Statement

All experimental materials described in this paper are available from the corresponding author on request. The protein structure coordinates and data have been deposited in the Protein Data Bank with deposition code 9R8S.

## Acknowledgements

We thank members of the Hagelueken group (University of Bonn) for sharing data prior to publication and for helpful discussions. We thank Shirley Graham and Sabine Grüschow for technical advice and help and the St Andrews Mass Spectrometry Unit for expert technical assistance.

---

**Supplementary Table 1:** Data collection and refinement statistics for AcrIII-2.

|  |  |
| --- | --- |
| **Data Collection** |  |
| Space group | P3121 |
| Cell dimensions |  |
| a, b, c (Å) | 110.6, 110.6, 113.0 |
| $\alpha$, $\beta$, $\gamma$ (°) | 90, 90, 120 |
| Resolution (Å)<br>(high resolution) | 95.79 – 1.79 (1.82 – 1.79) |
| $R_{\text{merge}}$ | 0.115 (4.572) |
| $I/\sigma(I)$ | 17.4 (0.4) |
| Completeness (%) | 100 (100) |
| Aver. Redundancy | 38.9 (29.6) |
| CC half | 1.000 (0.345) |
| V<sub>m</sub> (Å<sup>3</sup>/Da) | 3.71 |
| Solvent (%) | 66.9 |
| **Refinement** |  |
| Unique reflections | 75593 (3754) |
| R<sub>work</sub> / R<sub>free</sub> | 16.5 / 20.7 |
| Geometric deviations |  |
| Bonds (Å) / Angles (°) | 0.005 / 1.314 |
| No. atoms (non H) |  |
| Protein | 3811 |
| Water | 204 |
| Phosphate | 10 |
| B factors (Å<sup>2</sup>) |  |
| Protein | 43.7 |
| Water | 44.1 |
| Phosphate | 46.2 |
| Ramachandran |  |
| Favored / outlier (%) | 96.8 / 0.21 |
| Molprobity score / centile | 1.37 / 97 |

\* Values in parentheses are for the highest-resolution shell.

**Supplementary Figure 1.**
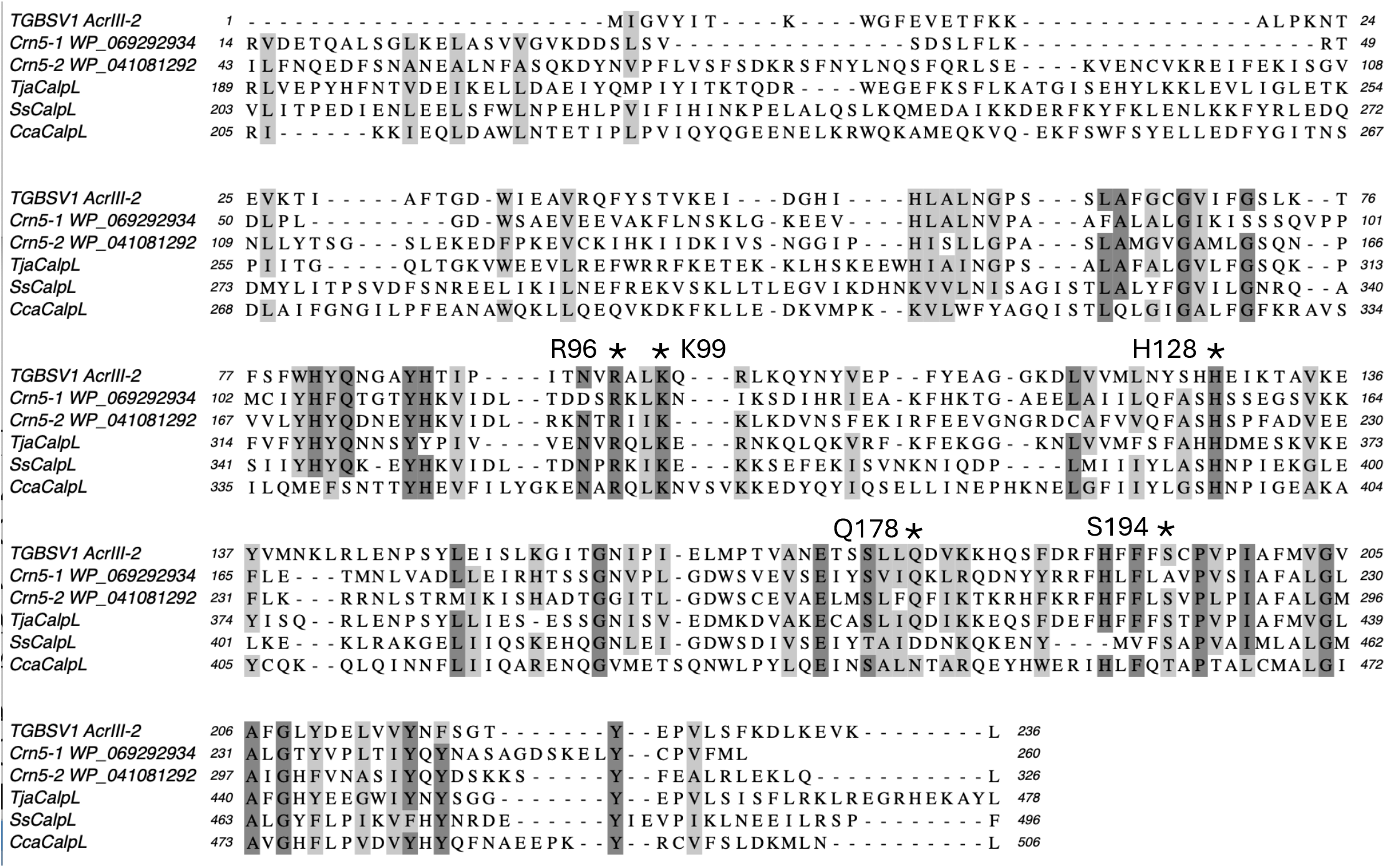
Sequence alignment of the TGBSV SAVED domain protein AcrIII-2 with bacterial homologues. The CalpL proteins from *Thermocrinis jamiesonii* (Tja), *Sulfurihydrogenibium* spp (Ss) and*Candidatus* Cloacimonas acidaminovorans (Cca) are shown. Crn5-1 and 5-2 are presumed CRISPR ring nucleases. Residues investigated by site directed mutagenesis are labelled.

**Supplementary Figure 2.**
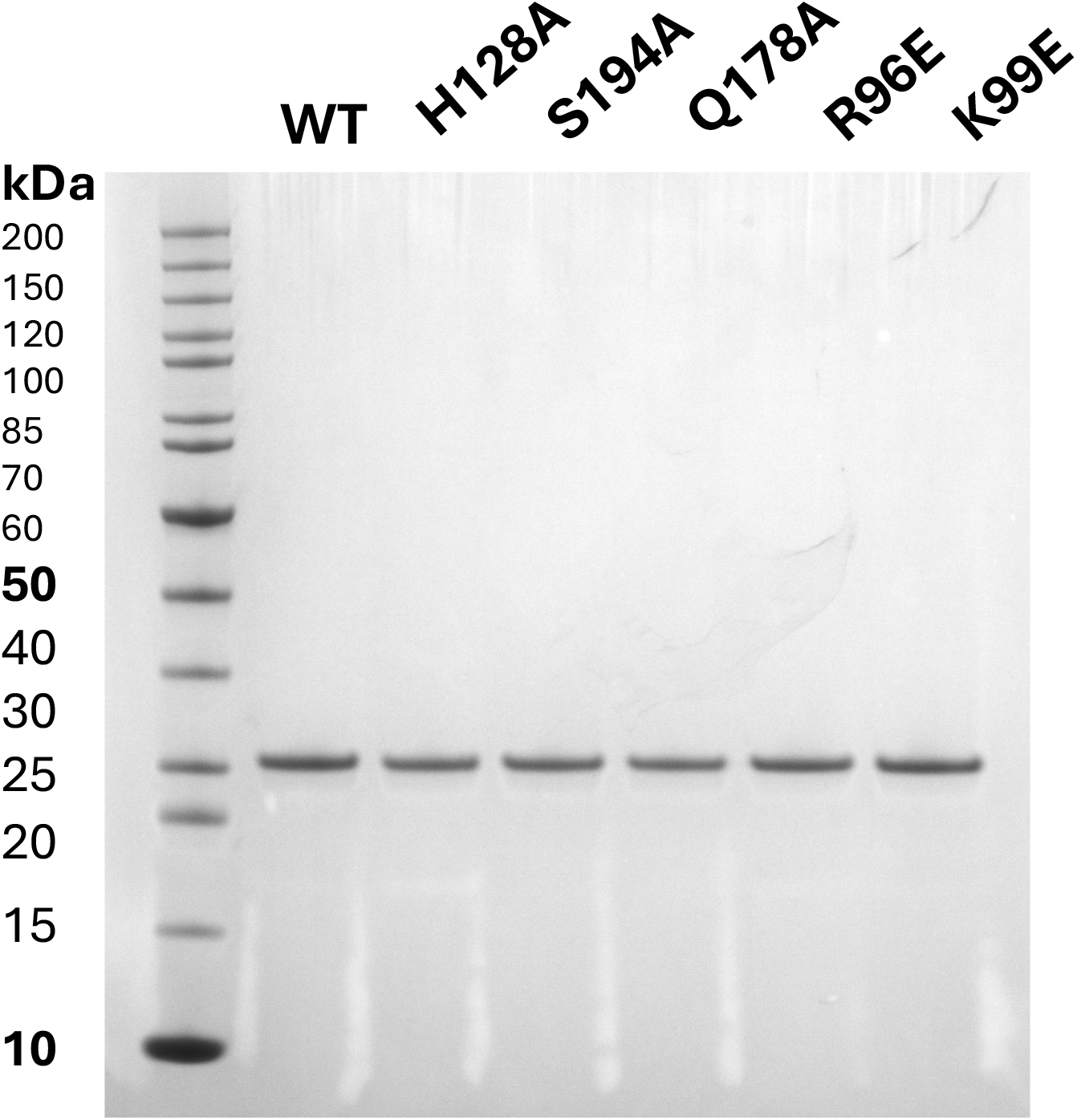
Purified AcrIII-2 wild-type and variant proteins.

**Supplementary Figure 3.**
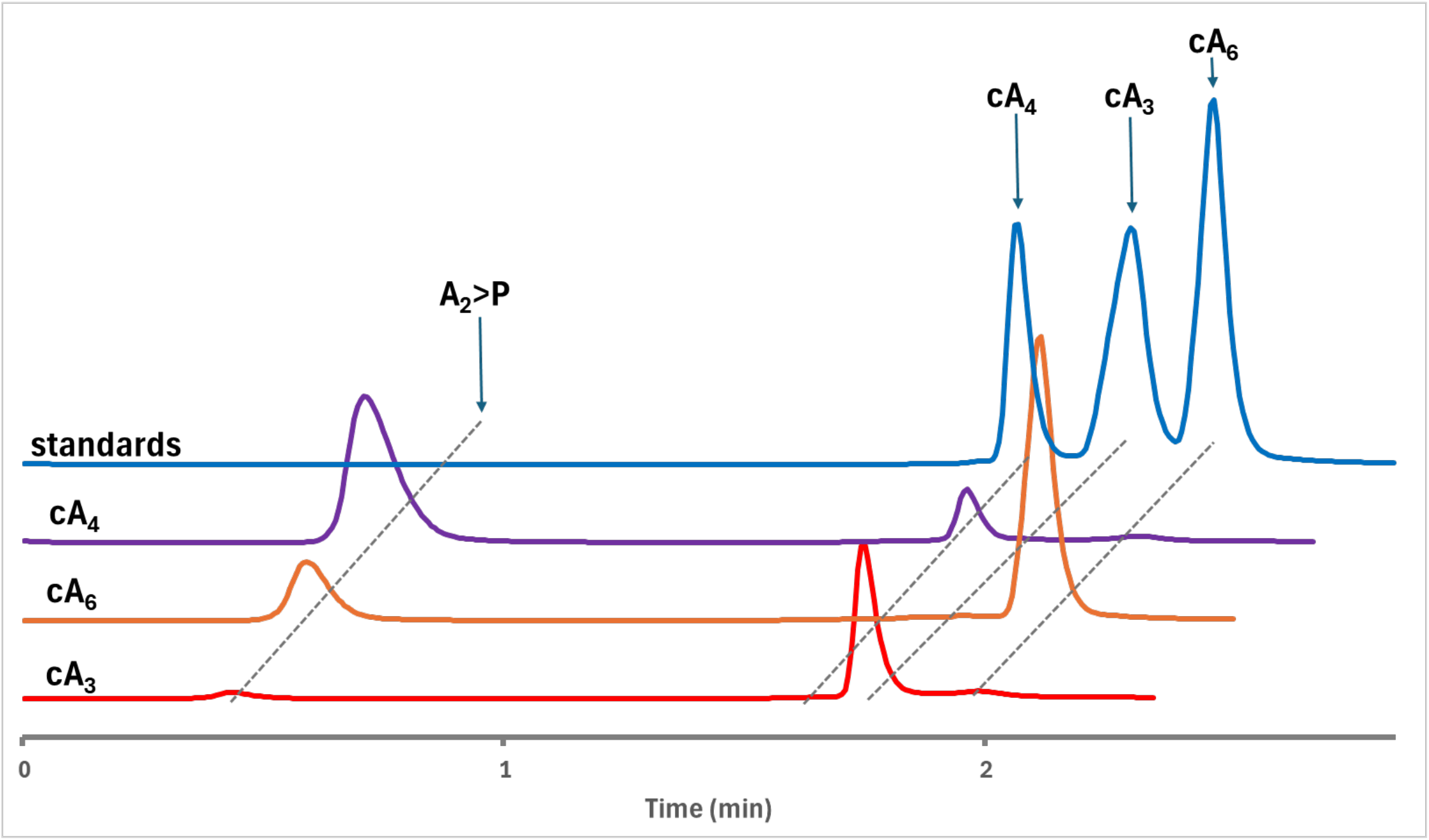
AcrIII-2 is cA_4_specific. AcrIII-2 was incubated with cA_3_, cA_4_, and cA_6_, respectively for 30 min at 60 °C. AcrIII-2 primarily degrades cA_4_, although cA_6_ was degraded to a lesser extent. AcrIII-2 showed minimal ring nuclease activity with cA_3_.

**Supplementary Figure 4.**
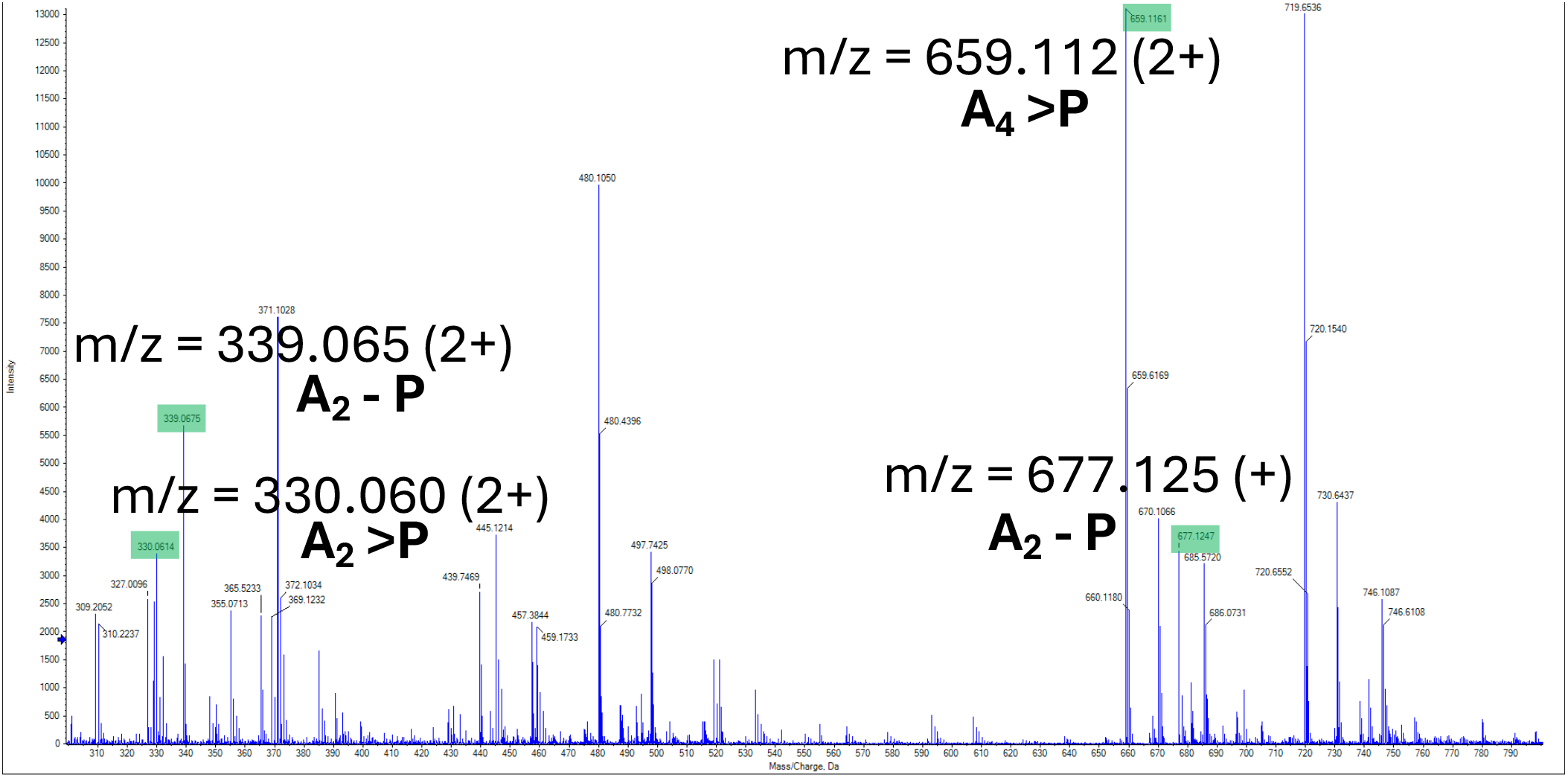
Mass spectrometry analysis of the initial product of cA_4_ degradation by AcrIII-2. (rightmost peak in Figure 1). This identifies A_4_>P as the most likely initial product, as observed for the CalpL proteins.

**Supplementary Figure 5.**
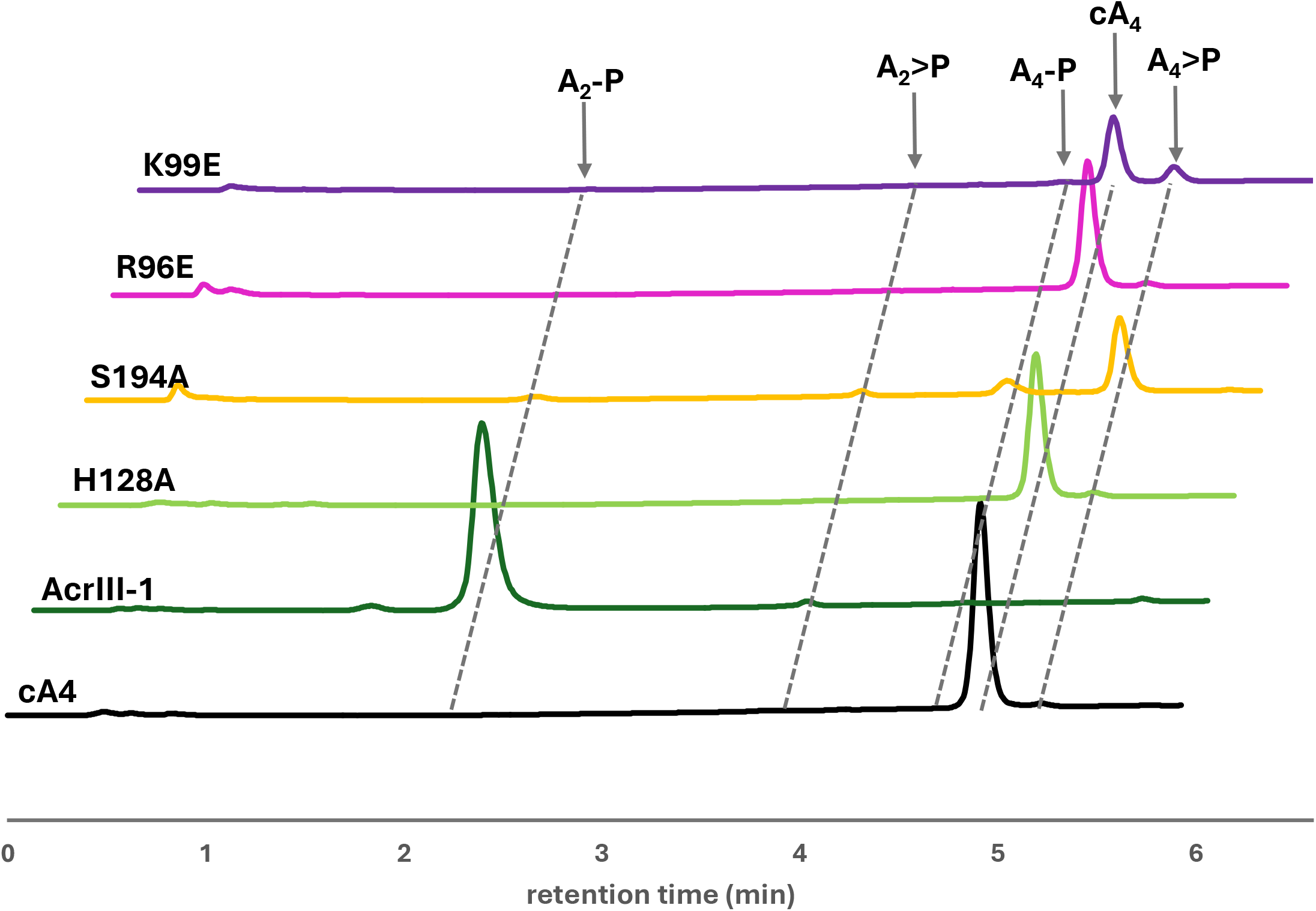
Activity of mutant AcrIII-2 at 1:1 cA_4_ to AcrIII-2. While H128A and R96E were still completely catalytically dead. K99E generated small amounts of A_4_>P, and S194A completely converted cA_4_ to linear tetranucleotide products.

## References

1 Millman, A., Melamed, S., Leavitt, A., Doron, S., Bernheim, A., Hor, J., Garb, J., Bechon, N., Brandis, A., Lopatina, A., Ofir, G., Hochhauser, D., Stokar-Avihail, A., Tal, N., Sharir, S., Voichek, M., Erez, Z., Ferrer, J. L. M., Dar, D., Kacen, A., Amitai, G. and Sorek, R. (2022) An expanded arsenal of immune systems that protect bacteria from phages. Cell host & microbe. 30, 1556–1569 e1555

2 Tesson, F., Herve, A., Mordret, E., Touchon, M., d’Humieres, C., Cury, J. and Bernheim, A. (2022) Systematic and quantitative view of the antiviral arsenal of prokaryotes. Nat Commun. 13, 2561

3 Payne, L. J., Meaden, S., Mestre, M. R., Palmer, C., Toro, N., Fineran, P. C. and Jackson, S. A. (2022) PADLOC: a web server for the identification of antiviral defence systems in microbial genomes. Nucleic Acids Res. 50, W541–W550

4 Sun, L., Wu, J., Du, F., Chen, X. and Chen, Z. J. (2013) Cyclic GMP-AMP synthase is a cytosolic DNA sensor that activates the type I interferon pathway. Science. 339, 786–791

5 Lowey, B., Whiteley, A. T., Keszei, A. F. A., Morehouse, B. R., Mathews, I. T., Antine, S. P., Cabrera, V. J., Kashin, D., Niemann, P., Jain, M., Schwede, F., Mekalanos, J. J., Shao, S., Lee, A. S. Y. and Kranzusch, P. J. (2020) CBASS Immunity Uses CARF-Related Effectors to Sense 3’-5’- and 2’-5’-Linked Cyclic Oligonucleotide Signals and Protect Bacteria from Phage Infection. Cell. 182, 38–49 e17

6 Duncan-Lowey, B., McNamara-Bordewick, N. K., Tal, N., Sorek, R. and Kranzusch, P. J. (2021) Effector-mediated membrane disruption controls cell death in CBASS antiphage defense. Mol Cell. 81, 5039–5051

7 Govande, A. A., Duncan-Lowey, B., Eaglesham, J. B., Whiteley, A. T. and Kranzusch, P. J. (2021) Molecular basis of CD-NTase nucleotide selection in CBASS anti-phage defense. Cell reports. 35, 109206

8 Niewoehner, O., Garcia-Doval, C., Rostol, J. T., Berk, C., Schwede, F., Bigler, L., Hall, J., Marraffini, L. A. and Jinek, M. (2017) Type III CRISPR-Cas systems produce cyclic oligoadenylate second messengers. Nature. 548, 543–548

9 Kazlauskiene, M., Kostiuk, G., Venclovas, C., Tamulaitis, G. and Siksnys, V. (2017) A cyclic oligonucleotide signaling pathway in type III CRISPR-Cas systems. Science. 357, 605–609

10 Hoikkala, V., Graham, S. and White, M. F. (2024) Bioinformatic analysis of type III CRISPR systems reveals key properties and new effector families. Nucleic Acids Res. 52, 7129–7141

11 Makarova, K. S., Timinskas, A., Wolf, Y. I., Gussow, A. B., Siksnys, V., Venclovas, C. and Koonin, E. V. (2020) Evolutionary and functional classification of the CARF domain superfamily, key sensors in prokaryotic antivirus defense. Nucleic Acids Res. 48, 8828–8847

12 Athukoralage, J. S., Graham, S., Rouillon, C., Grüschow, S., Czekster, C. M. and White, M. F. (2020) The dynamic interplay of host and viral enzymes in type III CRISPR-mediated cyclic nucleotide signalling. eLife. 9, e55852

13 Rouillon, C., Athukoralage, J. S., Graham, S., Gruschow, S. and White, M. F. (2018) Control of cyclic oligoadenylate synthesis in a type III CRISPR system. eLife. 7, e36734

14 Athukoralage, J. S., Rouillon, C., Graham, S., Grüschow, S. and White, M. F. (2018) Ring nucleases deactivate Type III CRISPR ribonucleases by degrading cyclic oligoadenylate. Nature. 562, 277–280

15 Hoikkala, V., Chi, H., Grüschow, S., Graham, S. and White, M. F. (2024) Diversity and abundance of ring nucleases in type III CRISPR-Cas loci. bioRxiv, 2024.2009.2024.614671

16 Athukoralage, J. S., McMahon, S. A., Zhang, C., Gruschow, S., Graham, S., Krupovic, M., Whitaker, R. J., Gloster, T. M. and White, M. F. (2020) An anti-CRISPR viral ring nuclease subverts type III CRISPR immunity. Nature. 577, 572–575

17 Hobbs, S. J., Wein, T., Lu, A., Morehouse, B. R., Schnabel, J., Leavitt, A., Yirmiya, E., Sorek, R. and Kranzusch, P. J. (2022) Phage anti-CBASS and anti-Pycsar nucleases subvert bacterial immunity. Nature. 605, 522–526

18 Cao, X., Xiao, Y., Huiting, E., Cao, X., Li, D., Ren, J., Fedorova, I., Wang, H., Guan, L., Wang, Y., Li, L., Bondy-Denomy, J. and Feng, Y. (2023) Phage anti-CBASS protein simultaneously sequesters cyclic trinucleotides and dinucleotides. Mol Cell 84, 375–385.

19 Cook, R., Brown, N., Redgwell, T., Rihtman, B., Barnes, M., Clokie, M., Stekel, D. J., Hobman, J., Jones, M. A. and Millard, A. (2021) INfrastructure for a PHAge REference Database: Identification of Large-Scale Biases in the Current Collection of Cultured Phage Genomes. Phage (New Rochelle). 2, 214–223

20 Palmer, M., Hedlund, B. P., Roux, S., Tsourkas, P. K., Doss, R. K., Stamereilers, C., Mehta, A., Dodsworth, J. A., Lodes, M., Monsma, S., Glavina Del Rio, T., Schoenfeld, T. W., Eloe-Fadrosh, E. A. and Mead, D. A. (2020) Diversity and Distribution of a Novel Genus of Hyperthermophilic Aquificae Viruses Encoding a Proof-Reading Family-A DNA Polymerase. Front Microbiol. 11, 583361

21 Rouillon, C., Schneberger, N., Chi, H., Blumenstock, K., Da Vela, S., Ackermann, K., Moecking, J., Peter, M. F., Boenigk, W., Seifert, R., Bode, B. E., Schmid-Burgk, J. L., Svergun, D., Geyer, M., White, M. F. and Hagelueken, G. (2023) Antiviral signalling by a cyclic nucleotide activated CRISPR protease. Nature. 614, 168–174

22 Binder, S. C., Schneberger, N., Schmitz, M., Engeser, M., Geyer, M., Rouillon, C. and Hagelueken, G. (2024) The SAVED domain of the type III CRISPR protease CalpL is a ring nuclease. Nucleic Acids Res. 52, 10520–10532

23 Smalakyte, D., Ruksenaite, A., Sasnauskas, G., Tamulaitiene, G. and Tamulaitis, G. (2024) Filament formation activates protease and ring nuclease activities of CRISPR Lon-SAVED. Mol Cell 84, 4239–4255.

24 Athukoralage, J. S., McQuarrie, S., Gruschow, S., Graham, S., Gloster, T. M. and White, M. F. (2020) Tetramerisation of the CRISPR ring nuclease Crn3/Csx3 facilitates cyclic oligoadenylate cleavage. eLife. 9, e57627

25 Brown, S., Gauvin, C. C., Charbonneau, A. A., Burman, N. and Lawrence, C. M. (2020) Csx3 is a cyclic oligonucleotide phosphodiesterase associated with type III CRISPR-Cas that degrades the second messenger cA(4). J Biol Chem. 295, 14963–14972

26 Athukoralage, J. S., Graham, S., Gruschow, S., Rouillon, C. and White, M. F. (2019) A Type III CRISPR Ancillary Ribonuclease Degrades Its Cyclic Oligoadenylate Activator. J Mol Biol. 431, 2894–2899

27 Wente, S. R. and Schachman, H. K. (1987) Shared active sites in oligomeric enzymes: model studies with defective mutants of aspartate transcarbamoylase produced by site-directed mutagenesis. Proc Natl Acad Sci U S A. 84, 31–35

28 McMahon, S. A., Zhu, W., Graham, S., Rambo, R., White, M. F. and Gloster, T. M. (2020) Structure and mechanism of a Type III CRISPR defence DNA nuclease activated by cyclic oligoadenylate. Nat Commun. 11, 500

29 Zhu, W., McQuarrie, S., Gruschow, S., McMahon, S. A., Graham, S., Gloster, T. M. and White, M. F. (2021) The CRISPR ancillary effector Can2 is a dual-specificity nuclease potentiating type III CRISPR defence. Nucleic Acids Res. 49, 2777–2789

30 Rostol, J. T., Xie, W., Kuryavyi, V., Maguin, P., Kao, K., Froom, R., Patel, D. J. and Marraffini, L. A. (2021) The Card1 nuclease provides defence during type-III CRISPR immunity. Nature. 590, 614–629

31 Baca, C. F., Yu, Y., Rostol, J. T., Majumder, P., Patel, D. J. and Marraffini, L. A. (2024) The CRISPR effector Cam1 mediates membrane depolarization for phage defence. Nature. 625, 797–804

32 Rouillon, C., Athukoralage, J. S., Graham, S., Grüschow, S. and White, M. F. (2019) Investigation of the cyclic oligoadenylate signalling pathway of type III CRISPR systems. Methods Enzymol. 616, 191–218

33 Winter, G. (2010) xia2: an expert system for macromolecular crystallography data reduction. J Appl Crystallogr. 43, 186–190

34 McCoy, A. J., Grosse-Kunstleve, R. W., Adams, P. D., Winn, M. D., Storoni, L. C. and Read, R. J. (2007) Phaser crystallographic software. J Appl Crystallogr. 40, 658–674

35 Winn, M. D., Ballard, C. C., Cowtan, K. D., Dodson, E. J., Emsley, P., Evans, P. R., Keegan, R. M., Krissinel, E. B., Leslie, A. G., McCoy, A., McNicholas, S. J., Murshudov, G. N., Pannu, N. S., Potterton, E. A., Powell, H. R., Read, R. J., Vagin, A. and Wilson, K. S. (2011) Overview of the CCP4 suite and current developments. Acta Crystallogr D Biol Crystallogr. 67, 235–242

36 Jumper, J., Evans, R., Pritzel, A., Green, T., Figurnov, M., Ronneberger, O., Tunyasuvunakool, K., Bates, R., Zidek, A., Potapenko, A., Bridgland, A., Meyer, C., Kohl, S. A. A., Ballard, A. J., Cowie, A., Romera-Paredes, B., Nikolov, S., Jain, R., Adler, J., Back, T., Petersen, S., Reiman, D., Clancy, E., Zielinski, M., Steinegger, M., Pacholska, M., Berghammer, T., Bodenstein, S., Silver, D., Vinyals, O., Senior, A. W., Kavukcuoglu, K., Kohli, P. and Hassabis, D. (2021) Highly accurate protein structure prediction with AlphaFold. Nature. 596, 583–589

37 Mirdita, M., Schutze, K., Moriwaki, Y., Heo, L., Ovchinnikov, S. and Steinegger, M. (2022) ColabFold: making protein folding accessible to all. Nat Methods. 19, 679–682

38 Liebschner, D., Afonine, P. V., Baker, M. L., Bunkoczi, G., Chen, V. B., Croll, T. I., Hintze, B., Hung, L. W., Jain, S., McCoy, A. J., Moriarty, N. W., Oeffner, R. D., Poon, B. K., Prisant, M. G., Read, R. J., Richardson, J. S., Richardson, D. C., Sammito, M. D., Sobolev, O. V., Stockwell, D. H., Terwilliger, T. C., Urzhumtsev, A. G., Videau, L. L., Williams, C. J. and Adams, P. D. (2019) Macromolecular structure determination using X-rays, neutrons and electrons: recent developments in Phenix. Acta Crystallogr D Struct Biol. 75, 861–877

39 Murshudov, G. N., Skubak, P., Lebedev, A. A., Pannu, N. S., Steiner, R. A., Nicholls, R. A., Winn, M. D., Long, F. and Vagin, A. A. (2011) REFMAC5 for the refinement of macromolecular crystal structures. Acta Crystallogr D. 67, 355–367

40 Casanal, A., Lohkamp, B. and Emsley, P. (2020) Current developments in Coot for macromolecular model building of Electron Cryo-microscopy and Crystallographic Data. Protein Sci. 29, 1069–1078

41 Chen, V. B., Arendall, W. B., 3rd, Headd, J.J., Keedy, D. A., Immormino, R. M., Kapral, G. J., Murray, L. W., Richardson, J. S. and Richardson, D. C. (2010) MolProbity: all-atom structure validation for macromolecular crystallography. Acta Crystallogr D Biol Crystallogr. 66, 12–21

